# Neuroticism alters the transcriptome of the frontal cortex to contribute to the cognitive decline and onset of Alzheimer’s disease

**DOI:** 10.1101/2020.08.21.261677

**Authors:** Céline H. De Jager, Charles C. White, David A. Bennett, Yiyi Ma

**Author notes:** Corresponding author: Yiyi Ma, M.D., Ph.D., Center for Translational & Computational Neuroimmunology, Department of Neurology, Columbia University Medical Center, 630 West 168.

## Abstract

Accumulating evidence has suggested that the molecular transcriptional mechanism contributes to Alzheimer’s disease (AD) and its endophenotypes of cognitive decline and neuropathological traits, β-amyloid (Aβ) and phosphorylated tangles (TAU). However, it is unknown what is the impact of the AD risk factors, personality characteristics assessed by the NEO Five-Factor Inventory, on the human brain’s transcriptome. Using postmortem human brain samples from 466 subjects, we found that neuroticism has a significant overall impact on the brain transcriptome (omnibus *P*=0.005) but not the other 4 personality characteristics. Focused on those cognitive decline related gene co-expressed modules, neuroticism has nominally significant associations (*P*<0.05) with four neuronal modules, which are more related to PHFtau than Aβ across all eight brain regions. Furthermore, the effect of neuroticism on cognitive decline and AD might be mediated through the expression of module 7 and TAU pathology (*P*=0.008). To conclude, neuroticism has overall impact on human brains’ transcriptome and its effect on cognitive decline and AD might be mediated through TAU pathology related gene transcription mechanism.

## Introduction

Alzheimer’s disease (AD) is the most common form of aging-related dementia, affecting over 5.8 million people in the United States alone (www.alz.org). Many risk factors that contribute to AD risk have been identified; some are genetic while others are environmental exposures or life experiences. For some risk factors, e.g., *APOE ε4,* the series of molecular events that lead to the accumulation of β-amyloid (Aβ) and phosphorylated tangles (TAU), subsequent loss of cognitive function, and, eventually, a diagnosis of dementia has been well documented.^1^ However, other risk factors remain poorly characterized at the molecular level. Here, we explored the molecular mechanism linking neuroticism to AD risk.^2^ Understanding this mechanism may yield important clues to the changes in RNA expression that contribute to the onset of AD and new avenues for developing therapeutics. Neuroticism, or proneness to distress, – is associated with faster cognitive decline and greater AD risk^3^. By contrast, conscientiousness – being organized, completing purposeful action, and having a drive to achieve – is associated with slower cognitive decline and lower AD risk^3^. While both have previously been reported to contribute to AD^4, 5^, the role of neuroticism has been more consistently replicated.^3^ The molecular composition of these personality traits and the series of molecular events that lead from personality traits to AD in the brain is unknown.

Recently, large-scale molecular data – such as RNA sequence (RNAseq) data from the dorsolateral prefrontal cortex (DLPFC)^6^ – have been generated from the brains of individuals who were deeply characterized while they were alive. Specifically, we repurposed data^6^ generated from 542 participants in two cohort studies of cognitive aging, the Religious Orders Study (ROS) and the Rush Memory and Aging Project (MAP), which include prospective brain autopsy of each participant.^7^ We previously examined the molecular features of the aging cortex that relate to AD pathology and cognitive decline.^6^ Here, we investigate the molecular consequences of personality traits leveraging data from the NEO Five Factor personality inventory. The DLPFC is one brain region that has been implicated in neuroticism based on magnetic resonance imaging (MRI).^8^

## Materials and Methods

### Participants

The Religious Orders Study (ROS) and the Rush Memory and Aging Project (MAP) are longitudinal studies which have enrolled >3,600 subjects without known dementia at baseline.^7^ Both studies were approved by an Institutional Review Board of Rush University Medical Center. All participants signed an informed consent, an Anatomical Gift Act, and a repository consent allowing their data to be repurposed. The participants undergo detailed cognitive testing on an annual basis and other ante-mortem phenotyping including the five personal traits: neuroticism, conscientiousness, extraversion, openness, and agreeableness. These personal variables are measured using 12 items from the NEO Five-Factor Inventory, and each item has scores which range from 0 to 4 and are summed to yield a maximum composite score of 48, where a higher score indicates greater magnitude of these personal traits ^5, 9–11^. The two studies are run by the same group of investigators and are designed to be analyzed jointly as the same procedures are used to capture these traits in both studies.

After death, a structured neuropathologic examination is performed to obtain quantitative measures of Aβ, TAU, and other pathologies as well as a pathologic diagnosis of AD. Aβ protein is identified by molecularly-specific immunohistochemistry and quantified by image analysis ^12, 13^. TAU pathology is measured by immunohistochemistry using the AT8 antibody which recognizes a phosphprylated Tau peptide. The values are the percentage area occupied by Aβ using image analysis. TAU proteins represent the density using stereology. Both Aβ and TAU are measured in 8 human brain regions: hippocampus, entorhinal cortex, midfrontal cortex, inferior temporal, angular gyrus, calcarine cortex, anterior cingulate cortex, and superior frontal cortex ^12, 13^, and the mean across the 8 brain regions are used as the overall level of Aβ or TAU. Results are transformed by taking the square root to fit normal distribution. The current analyses was approved by the Institutional Review Board of Columbia University Irving Medical Center.

One hemisphere is frozen at the time of autopsy, and this material was accessed to generate RNAseq data from the DLPFC.

### Gene expression modules

For the current study, there are 466 subjects with both measurements of the brain transcriptome and at least one of the five personality traits. The detailed methods by which the RNAseq were generated and analyzed were described in a prior report ^6^. In brief, gray matter from the DLPFC were processed, and RNA was extracted from each sample (Qiagen’s miRNeasey mini kit and the RNase free DNAse Set). A cDNA library was prepared from each sample and then sequenced using the Illumina HiSeq platform with 101 bp paired-end reads and an average coverage of 50 million reads. The obtained sequences were aligned to the reference genome of GRCh37 using Bowtie ^14^, and the level of expression of individual genes were estimated by the RSEM v1.2.31 ^15^ package. This was followed by quantile normalization and batch effect removal using the Combat algorithm ^16^. At the quality control stage, those genes with less than 4 reads in 100 individuals are removed, leading to 13,484 genes with expression measures. Groups of co-expressed genes (“modules”) across the 542 subjects were defined using the SpeakEasy ^17^ consensus clustering algorithm; the 13,484 genes were collapsed into 47 modules of coexpressed genes with an average of 330 genes per module.

### Statistical analysis

We applied a generalized linear regression model using the R “glm” function to conduct the association tests in the study. These models adjust for the covariates of age at death, education, sex, race, study, and technical variables of gene expression experiments such as the experimental batches and RNA integrity score (RIN score). For the mediation analysis, we used the “mediation” R package (https://cran.rproject.org/web/packages/mediation/vignettes/mediation.pdf). We compared the mediation *P*-value to the direct effect *P*-value. A *P*-value of <0.05 is considered as statistical significance.

## Results

### Population characteristics

We first reproduced the role of neuroticism and conscientiousness in the subset of 466 deceased ROSMAP participants (mean of age at death = 88y and 62% female) which have (1) personality inventory data; (2) at least two measures of cognitive performance so that a person-specific slope of cognitive decline can be calculated and (3) transcriptomic data (**Table 1**). We found that neuroticism contributes to accelerated cognitive decline (beta=−2.11×10^−3^, *P*=3.48×10^−3^) while conscientiousness is protective against cognitive decline (beta=2.38×10^−3^, *P*=2.58×10^−2^) controlling for the age at death, sex, race, education and study center. The association with neuroticism remained significant after further controlling the neuropathologic indices of Aβ, Lewy body, and vascular risk factors. However, after controlling for TAU, the association of neuroticism and cognitive decline became borderline (beta=-1.24×10^−3^, *P*=5.3×10^−2^), which were even non-significant when the model adjust all the neuropathologic indices altogether (beta=-9.71×10^−4^, *P*=0.11) (**Supplementary table S1**). These results are very consistent with prior analyses^4, 5^ of subsets of the ROSMAP participants and their associated manuscripts which reported these two associations. Having established the role of these two personality traits in aging-related cognitive decline in our sample, we moved on to identify the molecular features which are associated with them.

**Table 1.** Demographic characteristics of the ROSMAP participants used in the analysis.

|  | Male | Female | All | *P* |
| --- | --- | --- | --- | --- |
| Number<sup>1</sup> | 179 | 287 | 466 |  |
| Age at death (y)<sup>2</sup> | 86 (6.6) | 90 (6.6) | 88 (6.8) | 2.56E-07 |
| Education (y)<sup>2</sup> | 17.5 (3.7) | 16.1 (3.2) | 16.7 (3.5) | 7.76E-05 |
| White (N, %)<sup>3</sup> | 176, 98% | 285, 99% | 461 (99%) | 0.59 |
| Neuroticism<sup>2</sup> | 16 (6.2) | 17 (6.7) | 17 (6.5) | 0.04 |
| Conscientiousness<sup>2</sup> | 33 (5.5) | 34 (4.6) | 34 (5.0) | 0.02 |
| Openness<sup>2</sup> | 25 (6.1) | 26 (4.8) | 25 (5.3) | 0.23 |
| Agreeableness<sup>2</sup> | 33 (4.4) | 34 (3.5) | 34 (3.9) | 0.08 |
| Extraversion<sup>2</sup> | 30 (5.9) | 30 (6.0) | 30 (6.0) | 0.27 |
| Aβ pathology (square root) | 1.3 (1.1) | 1.6 (1.1) | 1.5 (1.1) | 0.05 |
| TAU pathology (square root) | 1.8 (1.3) | 2.2 (1.2) | 2.0 (1.3) | 0.009 |

<sup>1</sup>N represent the total number of subjects with measurements of any of the five personality traits: neuroticism, conscientiousness, openness, agreeableness, and extraversion.
<sup>2</sup>represent the mean and standard deviation of each trait in all subjects, males and females and the *P* values of their differences between males and females using t test.
<sup>3</sup>represent the count and percentage of each trait in all subjects, males and females and the *P* values of their differences between males and females using chi-square test.
Abbreviations: Aβ , β-amyloid; TAU, abnormally phosphorylated Tau protein, AT8.

### Neuroticism has an overall impact on modules of co-expressed genes

We first assessed the impact of neuroticism on the aging DLPFC’s transcriptome by conducting an omnibus test, which evaluates the distribution of the associations with all the 47 cortical co-expressed gene modules as a whole. We find that neuroticism has a broad effect on cortical RNA profiles: there is an excess of suggestive associations that are unlikely to have occurred by chance (*P*=0.005) (**Fig. 1a**). However, none of the other four personality traits: conscientiousness, agreeableness, openness, extraversion, has an overall impact on the transcriptome (*P*>0.05) (**Supplementary Fig. S1**). Another way to illustrate this finding is to illustrate that, at a threshold of *P*<0.05, neuroticism is associated with 18 (38%) of the 47 cortical modules (**Fig. 1b**). For comparison, the other 4 personality traits altogether are only associated with 4 modules at this threshold (**Supplementary Fig. S1**). Conscientiousness influences 1 of these 4 modules, but this module is unrelated to cognitive decline. Thus, neuroticism has an outsized association with the transcriptome of the aging brain compared to other personality traits.

**Fig. 1.**
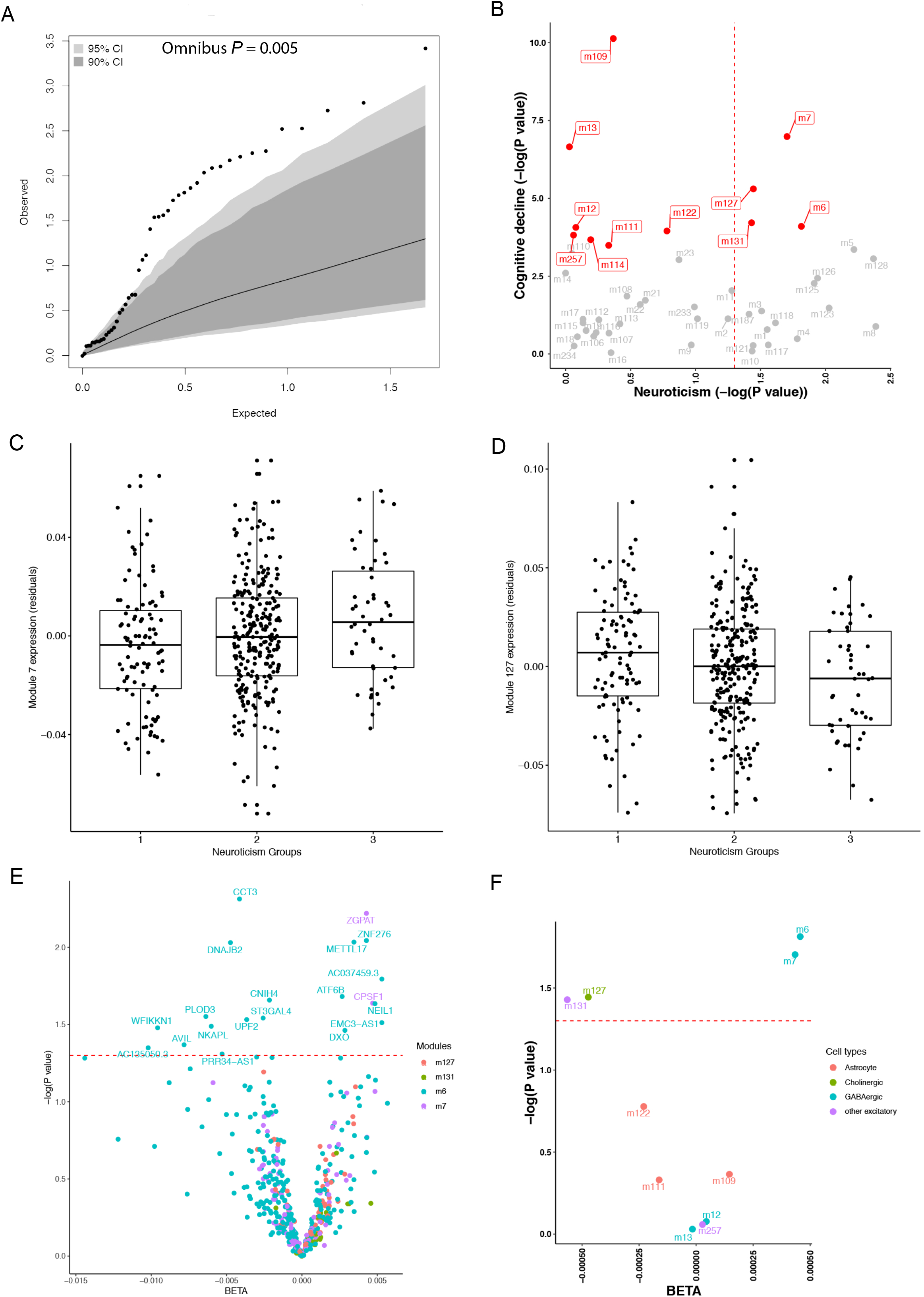
Neuroticism with overall or cognitive decline specific cortical co-expression gene modules. **a** The impact of neuroticism on the overall cortical co-expression gene modules. Each dot represents one module and its expected and observed associations with neuroticism (−log10 *P* values) are shown on X and Y axis, respectively. The expected *P* values assume a null distribution with no linear associations between neuroticism score and expression level of co-expressed gene module after adjusting age at death, sex, education, race, postmortem interval, study (ROS or MAP), RNA-seq batch and RIN score. The grey and dark areas indicate the extreme ranges of the QQ plot as generated by chance at a threshold of *P*=0.05 and *P*=0.10, respectively. The 95% and 90% confidence intervals were empirically derived by randomly assigning participants with the neuroticism score and repeating the analysis 1,000 times. Based on the distribution of the observed *P* values for all the 47 co-expressed gene modules, the overall association between neuroticism and module expression was unlikely to have occurred by chance (omnibus *P*=0.005). **b** The impact of neuroticism on the cognitive decline related co-expression gene modules. Each dot represents one module (the module number is listed next to each dot) and their previously reported *P* values for the cognitive decline (*Mostafavi 2018*) are shown on the Y axis and the P values (-log10 transformed) of the effect of neuroticism are shown on the X axis with the red dashed line for the threshold of *P*=0.05. The red dots are those that were previously reported to be associated with cognitive decline (*Mostafavi 2018*). **c** and **d** Distribution of cortical module m7 and m127 expression in individuals with different levels of neuroticism. Each dot represents one subject, and the subjects are distributed into 3 groups based on their neuroticism score: Group 1 with score from 0 to 12; Group 2 with score from 13 to 24; Group 3 with score from 25 to 36. On this scale, scores go from 0 (no neuroticism) to 36 (extensive neuroticism). **e** Volcano plot of the associations between neuroticism score and mRNA expression level of each of the 499 genes included in the top 4 modules associated with both neuroticism and cognitive decline with color codings of m7 in purple, m6 in blue, m127 in green, and m131 in red. **f** Volcano plot of the associations between neuroticism score and the 11 previously reported cognitive decline associated co-expressed gene modules with color codings of different cell type: astrocyte in red, cholinergic neuron in green, GABAergic neuron in blue, and other excitatory neuron in purple. For both volcano plots, X axis shows the BETA and Y axis shows the −log10 transformed *P* values with the red dashed line represents the threshold of *P*=0.05.

### Neuroticism has an impact on modules of co-expressed genes that are associated with cognitive decline

We previously reported 11 modules of co-expressed genes have significant associations with cognitive decline ^6^, and four of them reached nominal significance for the neuroticism, which are modules m7 (*P*=0.02), m6 (*P*=0.02), m127 (*P*=0.04), and m131 (*P*=0.04) (**Fig. 1b**). The association between neuroticism and the expression levels of each of the 4 modules are positive for m7 and m6 but negative for m127 and m131 (**Fig. 1c,d** & **Supplementary Fig. S2**). This means that subjects with more severe neuroticism syndrome have higher expression levels of the genes included in the m7 and m6 but lower expression levels of the genes included in the m127 and m131. We previously reported that subjects with accelerated cognitive decline have higher expression levels of genes in m7 and m6 but lower levels in m127 and m131 ^6^. In this case, the direction of the associations among neuroticism, modules and cognitive decline is consistent, showing that subjects with more severe neuroticism syndromes have more rapid cognitive decline, higher expression levels of m7 and m6 but lower levels of m127 and m131. Thus, we prioritized a set of modules that may be responding to greater neurotic behavior and cause downstream cognitive decline. However, none of the 11 modules display a relationship to conscientiousness, which is protective for AD; that process may work through other mechanisms or may work through another brain region.

We further conducted the association test with the individual gene included in the modules m7, m6, m127 and m131 to identify those specific genes related to neuroticism. Out of the 499 genes in these 4 modules, 20 have *P*<0.05, 18 of which are in m6 with 2 in m7 (**Fig. 1e**). The pathway enrichment analysis for these 20 genes shows in the involvement of mRNA surveillance pathway (FDR_*P*=0.008, hits=2 (2%)) (Data not shown).

We also annotated the cell types of the 4 modules associated with both neuroticism and cognitive decline (**Fig. 1f**). All of the 4 modules are annotated to be neurons. Both the modules m7 and m6 are enriched in the genes expressed by GABAergic neurons while the m127 is enriched in the genes expressed by cholinergic neurons and m131 is enriched in genes expressed by the other excitatory neurons.

### Neuroticism and cognitive decline associated co-expressed gene modules are related to AD pathology

We further analyzed which AD pathology, Aβ or TAU, are primarily involved in neuroticism pathology. Neuroticism has a positive association with TAU (*P*=0.03) but not Aβ (*P*=0.16) in this subsample of participants (**Fig. 2a**), both of which were the mean densities across 8 different brain regions. For the 4 modules associated with both neuroticism and cognitive decline, 3 of them have smaller *P* values for TAU than the *P* values for Aβ based on our previously reported associations ^6^ and only module m7 was reported to have significant association with the average level of TAU (**Fig. 2b**). More detailed analysis with the 8 region-specific measures of TAU and Aβ also suggest that the effect of module m7 on TAU outperformed all the other tested associations (**Fig. 2c**).

**Fig. 2.**
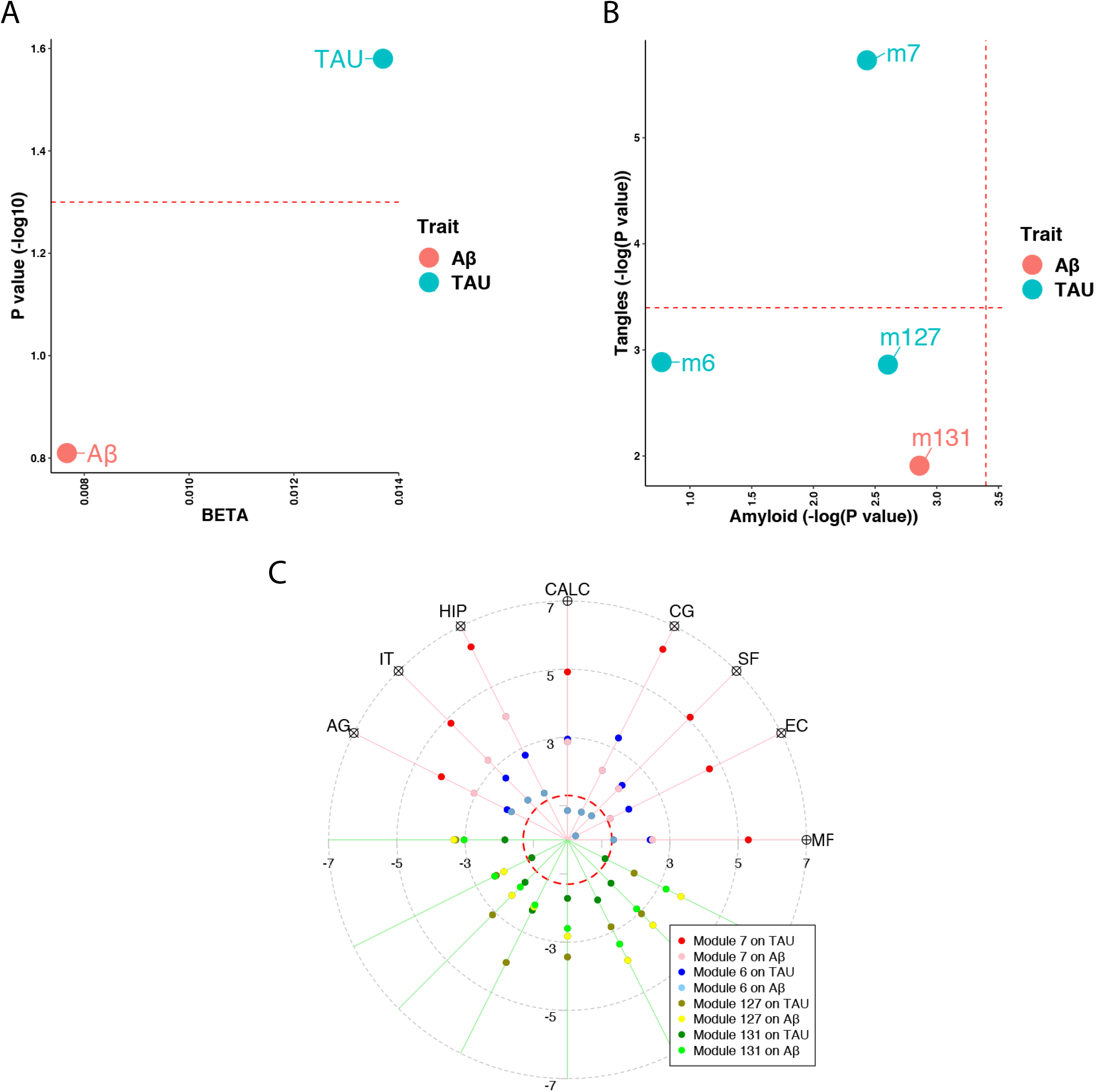
Neuroticism and cognitive decline associated co-expressed gene modules are more related to TAU than Aβ. **a** Effect of neuroticism on the average levels of brain TAU and Aβ, which were defined as the mean of the measurements across 8 different brain regions of midfrontal cortex (MF), entorhinal cortex (EC), superior frontal cortex (SF), anterior cingulate cortex (CG), calcarine cortex (CALC), hippocampus (HIP), inferior temporal cortex (IT), and angular gyrus (AG). X axis represent the BETAs and Y axis represent the −log10 transformed *P* values for the neuroticism score on either TAU (blue dot) or Aβ (red dot) after adjusting for the age at death, sex, postmortem interval, race, education, study (ROS or MAP). The horizontal red dashed line represents the threshold of *P*=0.05. **b** Comparisons of the associations to Aβ vs. TAU for the 4 modules related to both cognitive decline and neuroticism. Each dot represents one module with the *P* values (−log10 transformed) reported by Mostafavi 2018 for Aβ (X axis) and TAU (Y axis). Red dots show those modules with smaller reported P value for Aβ than TAU while blue dots show those modules with smaller reported P value for TAU than Aβ. **c** Target plot shows the associations between the 4 cognitive decline/neuroticism related modules and measures of Aβ and TAU across 8 different brain regions, which are displayed counter-clockwise from MF, EC, SF, CG, CALC, HIP, IT, and AG. Each dot on each axis represent the signed P value (-log10 transformed) of the association between each module and each measures of either Aβ or TAU at each brain region with the color codings of: module m7 on TAU in red and Aβ in pink, m6 on TAU in blue and Aβ in skyblue, m127 on TAU in gold and Aβ in yellow, and m131 on TAU in darker green and Aβ in green. The red dashed circle represents the threshold of *P*=0.05. The covariates include age at death, sex, postmortem interval, race, education, study (ROS or MAP), RIN and RNA experimental batch. Abbreviations: Aβ, β-amyloid; TAU, abnormally phosphorylated Tau protein, AT8.

### Mediation modeling to propose a sequence of events

Using a rigorous statistical methodology and our cross-sectional data obtained from autopsy material, we assessed the most likely location of each module in the sequence of events leading to AD to help to generate hypotheses that can be tested in preclinical and *in vitro* models. Mediation modeling suggests that the 4 modules of co-expressed genes which are altered in expression in the frontal cortex by neuroticism may contribute to accelerating aging-related cognitive decline. For example, using m7 which is the module most strongly associated with cognitive decline, neuroticism is associated with faster cognitive decline (*P*=0.008) through an association with higer module 7 than its direct causal effect on cognitive decline which is not mediated through m7 expression (*P*=0.042) (**Fig. 3 left**). We find similar results for modules m6 and m127 but not m131, which does not mediate the effect of neuroticism on TAU although TAU mediates the effect of m131 on cognitive decline (**Supplementary table S2**).

**Fig. 3.**
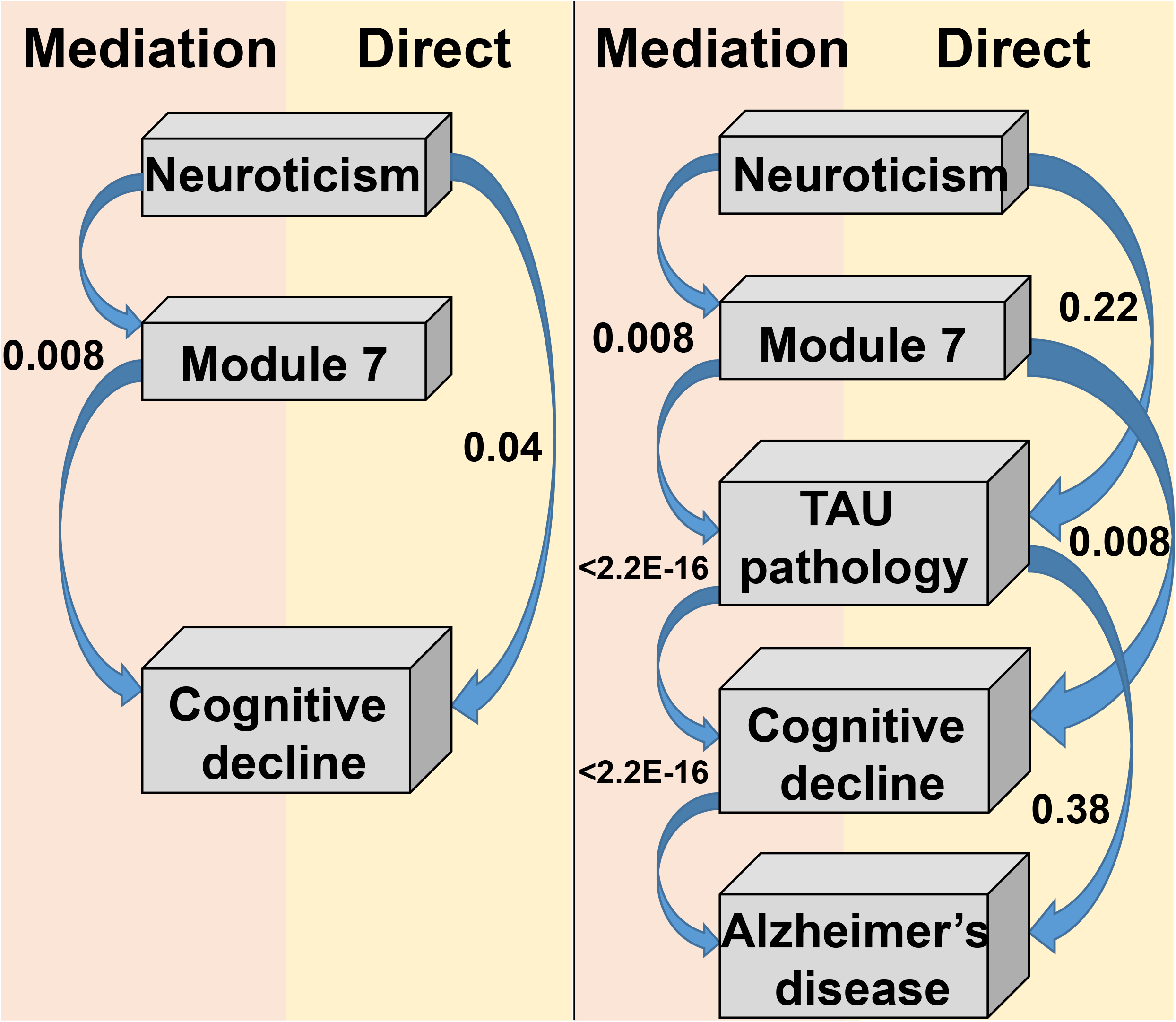
Module m7 mediates the effect of neuroticism on cognitive decline. **Left panel:** we present the most likely sequence of events based on our cross-sectional data. The most likely scenario is that being more neurotic leads to an increase in m7 which then contributes to an acceleration in cognitive decline. The p-value for this 0.008 while, in the same analysis, the direct effect of neuroticism on cognitive decline is much less (*P*=0.04). Not shown is an analysis which shows that m7 is much more likely to be downstream of neuroticism than upstream where an increase in m7 would make someone more neurotic. **Right panel:** we expand the model shown in the left panel with Tau pathology which is known to immediately precede cognitive decline and a diagnosis of AD which occurs after someone has begun to decline. Here, three mediation tests are presented: (1) Neuroticism→m7→Tau is more likely than a direct effect of Neuroticism→Tau, (2) m7→Tau→Cognitive decline is more likely than m7→Cognitive decline, and (3) Tau contributes to AD dementia by affecting the slope of cognitive decline (Tau→Cognitive decline→AD), which is the well-accepted model of AD. Abbreviations: TAU, abnormally phosphorylated Tau protein, AT8.

Based on our previous and current report that m7 is the module most directly contributing to the accumulation of TAU pathology ^6^, we added this variable to our model to more precisely evaluate the mechanism (**Fig. 3 right**). Thus, we propose that the most likely scenario in terms of the role of m7 in neuroticism’s effect on AD is the following sequence of events: more neuroticism → more m7 in frontal cortex → more TAU pathology accumulation → more cognitive decline → greater likelihood of developing AD dementia. m127 and m131 are not associated with TAU pathology and may therefore work through another mechanism to mediate the effects of neuroticism on cognitive decline (mediation p<0.01).

## Discussion

Neuroticism appears to have a broad impact on the biology of the older brain, as measured by RNA expression: approximately 1/5 of our modules of co-expressed genes are associated with an individual’s level of neuroticism at a nominal p<0.05 threshold. Thus, we have started to map out the molecular changes in the target organ (brain) that relate to neuroticism, an important personality trait that is a risk factor for AD and aging-related cognitive decline. The molecular substrate of neuroticism remains poorly understood as few studies have systematically assessed this question in large numbers of individuals, so this report is an important step forward in demonstrating a widespread biological effects of neuroticism in the frontal cortex. These large changes in older individuals beg the question of whether the changes are clinically meaningful: are they associated with symptoms, with changes in cognition?

We explored this question in detail and found that 3 of the modules associated with neuroticism are also associated with the slope of cognitive decline and a diagnosis of AD, providing a potential molecular link between AD and its risk factor, neuroticism. Since association studies of cross-sectional data, such as the one that we performed, cannot formally resolve the direction of causality, we used rigorous statistical methodology to test for mediation which enables us to propose a sequence of events that best fits our cross-sectional data. These analyses are very useful in generating hypotheses that can then be tested in cell culture or mouse model systems. In our analyses, we find that module m7, the module with the strongest association with neuroticism is most likely to increase in expression in response to increased neurotic behavior and to then contribute to cognitive decline (Fig. 3). These insights are important because, if validated in future studies, it could guide the development of future drugs to prevent AD. For example, a drug reducing m7 expression may be ineffective in preventing neuroticism, but it may be helpful to prevent the spread of negative molecular changes in the cortex that lead to cognitive decline and AD. Since personality traits like neuroticism cannot be readily changed, blocking the dysfunctional consequences of such personality traits may be the best option with which to manage individuals at risk of AD because of their personality traits.

We focused on neuroticism given that it is the personality trait most strongly associated with risk of AD and that its association has been well validated. Nonetheless, we also evaluated four other major personality traits in our analyses to be thorough, and, while we found some associations, they were much fewer than the neuroticism associations and did not relate to cognitive decline. It is possible that their effects may be more subtle and/or that we need to sample other brain regions to identify their effects.

Our study has many strengths, including its large sample size and largely community-based design, which facilitates repurposing our results and may offer insights translatable to the older community. We also have very deep RNAseq profiles, which enhanced the modules’ information content. Nonetheless, there are important limitations as well: we have already discussed the cross-sectional nature of the data that is a practical limitation of autopsy studies and prevents us from formally resolving the issue of causality. In addition, the average age at death was 88 years, which may make it challenging to extrapolate our results to the population of younger older individuals where many treatments for AD will need to be targeted.

Overall, we have prioritized a set of molecular changes in the frontal cortex that are involved in both neuroticism and aging-related cognitive decline/AD. We have also proposed a specific sequence of events that can now be tested in model systems. This hypothesis can be refined by performing many of the analyses reported here at the gene level instead of at the level of the module of co-expressed genes: this will facilitate validation studies and will be necessary to enable drug development efforts by presenting a specific target. We have therefore laid a strong foundation for future biological investigations of the effects of an important AD risk factor that affects many individuals: neuroticism.

## Acknowledgements

We are grateful to the participants in the Religious Order Study, the Rush Memory and Aging Project. This work is supported by the US National Institutes of Health U01AG61356, RP30AG10161, R01AG17917, R01AG15819. Ma Y. is supported by the 2019-AARF-644521.

## Author contributions

CHD and YM: study concept and design, data analysis and manuscript draft. CCW: guidance of omnibus test. DAB: provide data and critical review of manuscript.

## References

1. Theendakara V, Peters-Libeu CA, Bredesen DE, Rao RV. Transcriptional Effects of ApoE4: Relevance to Alzheimer’s Disease. Mol Neurobiol 2018; 55(6): 5243–5254.

2. Wilson RS, Begeny CT, Boyle PA, Schneider JA, Bennett DA. Vulnerability to stress, anxiety, and development of dementia in old age. Am J Geriatr Psychiatry 2011; 19(4): 327–334.

3. D’Iorio A, Garramone F, Piscopo F, Baiano C, Raimo S, Santangelo G. Meta-Analysis of Personality Traits in Alzheimer’s Disease: A Comparison with Healthy Subjects. J Alzheimers Dis 2018; 62(2): 773–787.

4. Wilson RS, Fleischman DA, Myers RA, Bennett DA, Bienias JL, Gilley DW et al. Premorbid proneness to distress and episodic memory impairment in Alzheimer’s disease. J Neurol Neurosurg Psychiatry 2004; 75(2): 191–195.

5. Wilson RS, Schneider JA, Arnold SE, Bienias JL, Bennett DA. Conscientiousness and the incidence of Alzheimer disease and mild cognitive impairment. Arch Gen Psychiatry 2007; 64(10): 1204–1212.

6. Mostafavi S, Gaiteri C, Sullivan SE, White CC, Tasaki S, Xu J et al. A molecular network of the aging human brain provides insights into the pathology and cognitive decline of Alzheimer’s disease. Nat Neurosci 2018; 21(6): 811–819.

7. Bennett DA, Buchman AS, Boyle PA, Barnes LL, Wilson RS, Schneider JA. Religious Orders Study and Rush Memory and Aging Project. J Alzheimers Dis 2018; 64(s1): S161–S189.

8. Lucas I, Balada F, Blanco E, Aluja A. Prefrontal cortex activity triggered by affective faces exposure and its relationship with neuroticism. Neuropsychologia 2019; 132: 107146.

9. Krueger KR, Wilson RS, Shah RC, Tang Y, Bennett DA. Personality and incident disability in older persons. Age Ageing 2006; 35(4): 428–433.

10. Wilson RS, Krueger KR, Gu L, Bienias JL, Mendes de Leon CF, Evans DA. Neuroticism, extraversion, and mortality in a defined population of older persons. Psychosom Med 2005; 67(6): 841–845.

11. Buchman AS, Boyle PA, Wilson RS, Leurgans SE, Arnold SE, Bennett DA. Neuroticism, extraversion, and motor function in community-dwelling older persons. Am J Geriatr Psychiatry 2013; 21(2): 145–154.

12. Bennett DA, Schneider JA, Arvanitakis Z, Wilson RS. Overview and findings from the religious orders study. Curr Alzheimer Res 2012; 9(6): 628–645.

13. Bennett DA, Schneider JA, Buchman AS, Barnes LL, Boyle PA, Wilson RS. Overview and findings from the rush Memory and Aging Project. Curr Alzheimer Res 2012; 9(6): 646–663.

14. Langmead B, Trapnell C, Pop M, Salzberg SL. Ultrafast and memory-efficient alignment of short DNA sequences to the human genome. Genome Biol 2009; 10(3): R25.

15. Li B, Dewey CN. RSEM: accurate transcript quantification from RNA-Seq data with or without a reference genome. BMC Bioinformatics 2011; 12: 323.

16. Johnson WE, Li C, Rabinovic A. Adjusting batch effects in microarray expression data using empirical Bayes methods. Biostatistics 2007; 8(1): 118–127.

17. Gaiteri C, Chen M, Szymanski B, Kuzmin K, Xie J, Lee C et al. Identifying robust communities and multi-community nodes by combining top-down and bottom-up approaches to clustering. Sci Rep 2015; 5: 16361.

