## Supplementary figures and images for "Neuroticism alters the transcriptome of the frontal cortex to contribute to the cognitive decline and onset of Alzheimer’s disease"

### Supplementary figure S1

A Conscienciousness

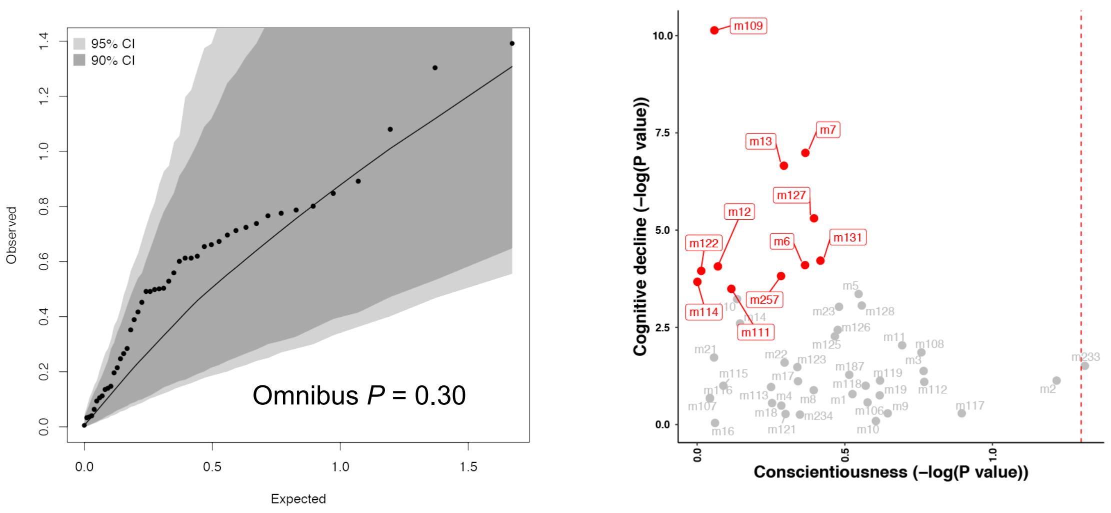


B Agreeableness

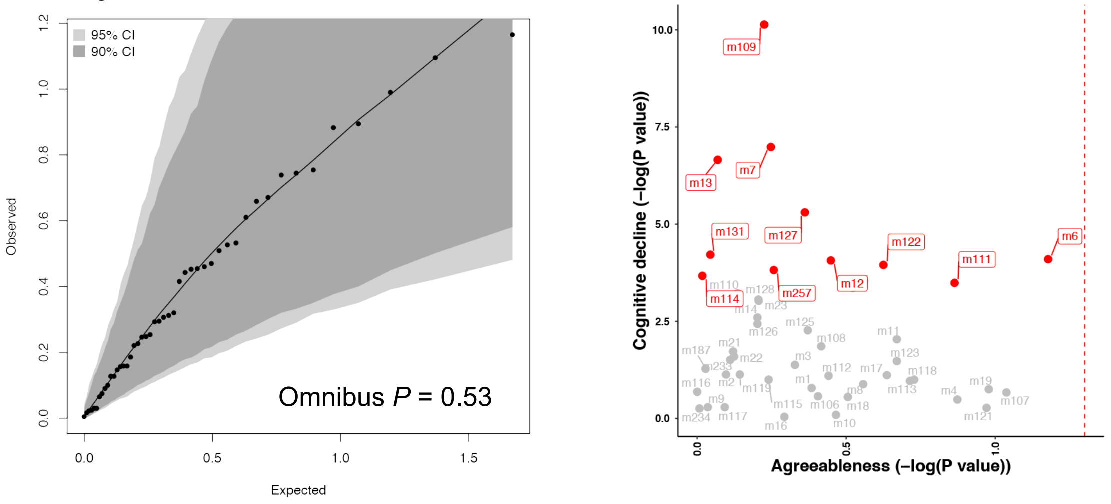


C Openness

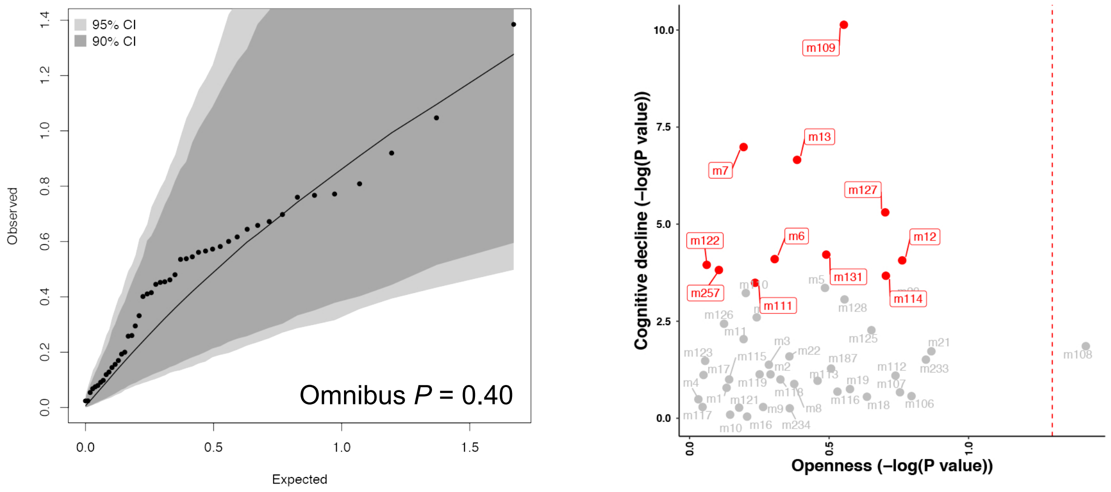


D Extraversion

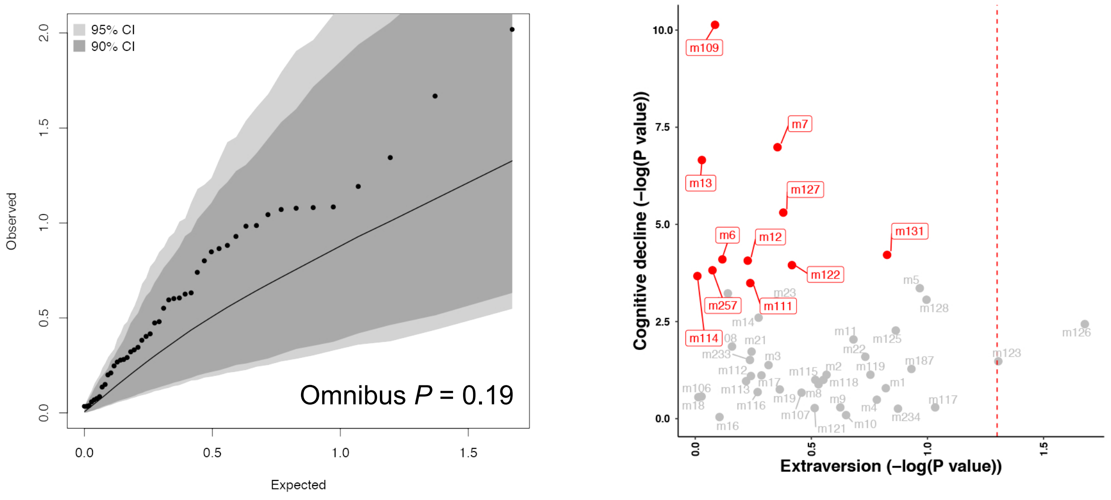

### Supplementary figure S2

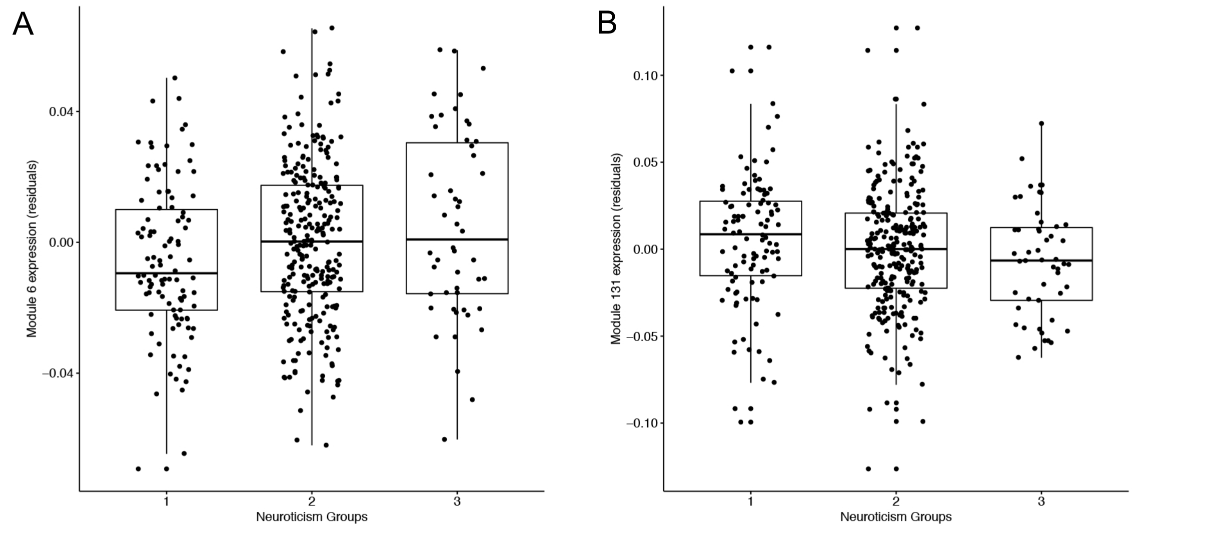
