## Supplementary File for "Neuroticism alters the transcriptome of the frontal cortex to contribute to the cognitive decline and onset of Alzheimer’s disease"

### SUPPLEMENTAL MATERIAL CONTENTS


#### SUPPLEMENTAL TABLES

**Supplementary table S1.** Confirmation of associations of neuroticism and cognitive decline within the subjects with measurements of postmortem brain gene expressions.
**Supplementary table S2.** Mediation analysis results.

#### SUPPLEMENTAL FIGURES

**Supplementary figure S1. The impact of other four personality traits on the overall cortical expressions of co-expressed genes**.
**Supplementary figure S2. Distribution of cortical module m6 and m131 expression in individuals with different levels of neuroticism**.

**Supplementary table S1.** Confirmation of associations of neuroticism and cognitive decline within the subjects with measurements of postmortem brain gene expressions.

| Model | Trait | N | BETA | SE | P |
|---|---|---|---|---|---|
| Model A | Neuroticism | 402 | -2.11E-03 | 7.19E-04 | 3.48E-03 |
| | Conscientiousness | 293 | 2.38E-03 | 1.06E-03 | 2.58E-02 |
| | Openness | 269 | 5.97E-04 | 1.14E-03 | 6.01E-01 |
| | Agreeableness | 270 | 2.74E-03 | 1.49E-03 | 6.74E-02 |
| | Extraversion | 441 | 3.19E-04 | 7.75E-04 | 6.81E-01 |
| Model A + Amyloid | Neuroticism | 397 | -1.71E-03 | 6.85E-04 | 1.28E-02 |
| | Conscientiousness | 288 | 1.56E-03 | 1.03E-03 | 1.32E-01 |
| | Openness | 264 | 6.78E-04 | 1.09E-03 | 5.36E-01 |
| | Agreeableness | 265 | 2.22E-03 | 1.42E-03 | 1.18E-01 |
| | Extraversion | 436 | 3.44E-04 | 7.36E-04 | 6.40E-01 |
| Model A + Tangles | Neuroticism | 397 | -1.24E-03 | 6.37E-04 | 5.30E-02 |
| | Conscientiousness | 288 | 1.32E-03 | 9.37E-04 | 1.59E-01 |
| | Openness | 264 | 1.51E-03 | 1.00E-03 | 1.31E-01 |
| | Agreeableness | 265 | 2.33E-03 | 1.29E-03 | 7.17E-02 |
| | Extraversion | 436 | 6.03E-04 | 6.64E-04 | 3.64E-01 |
| Model A + Lewy body | Neuroticism | 402 | -1.97E-03 | 7.05E-04 | 5.47E-03 |
| | Conscientiousness | 293 | 2.28E-03 | 1.07E-03 | 3.35E-02 |
| | Openness | 269 | 8.23E-04 | 1.15E-03 | 4.76E-01 |
| | Agreeableness | 270 | 2.84E-03 | 1.49E-03 | 5.79E-02 |
| | Extraversion | 441 | 4.06E-04 | 7.62E-04 | 5.94E-01 |
| Model A + Macro-infarcts | Neuroticism | 402 | -2.08E-03 | 7.14E-04 | 3.75E-03 |
| | Conscientiousness | 293 | 2.48E-03 | 1.06E-03 | 1.96E-02 |
| | Openness | 269 | 6.89E-04 | 1.15E-03 | 5.49E-01 |
| | Agreeableness | 270 | 2.85E-03 | 1.49E-03 | 5.66E-02 |
| | Extraversion | 441 | 2.87E-04 | 7.71E-04 | 7.10E-01 |
| Model A + Micro-infarcts | Neuroticism | 402 | -2.01E-03 | 7.18E-04 | 5.45E-03 |
| | Conscientiousness | 293 | 2.58E-03 | 1.06E-03 | 1.54E-02 |
| | Openness | 269 | 5.59E-04 | 1.15E-03 | 6.26E-01 |
| | Agreeableness | 270 | 3.21E-03 | 1.49E-03 | 3.19E-02 |
| | Extraversion | 441 | 2.87E-04 | 7.71E-04 | 7.10E-01 |
| Model A + All 5 neuropathologic indices | Neuroticism | 397 | -9.71E-04 | 6.11E-04 | 1.13E-01 |
| | Conscientiousness | 288 | 1.15E-03 | 9.11E-04 | 2.09E-01 |
| | Openness | 264 | 1.41E-03 | 9.69E-04 | 1.47E-01 |
| | Agreeableness | 265 | 2.73E-03 | 1.25E-03 | 2.99E-02 |
| | Extraversion | 436 | 6.56E-04 | 6.37E-04 | 3.04E-01 |

Note: N, BETA, SE, and P represent the analysis sample size, regression coefficient, standard error, and *P* value of the effect of each of the five personality trait on the cognitive decline.
The covariates in model A include age at death, sex, race, education, and study (not for openness and agreeableness). The model with the additional adjustment of the neuropathologic indices also include adjustment of postmortem interval.
Abbreviations: Aβ, β-amyloid; TAU, abnormally phosphorylated Tau protein, AT8.

**Supplementary table S2. Mediation analysis results.**

| Exposure | Mediator | Outcome | N | Analysis | Coef. [95% CI] | *P* |
|---|---|---|---|---|---|---|
| Neuroticism | m7 | TAU | 388 | ACME | 0.01 [0.006,0.01] | 0.008 |
| | | | | ADE | 0.01 [-0.01,0.03] | 0.224 |
| | | | | Proportion | 0.34 [-0.8,2.4] | 0.082 |
| | | | | Total | 0.02 [-0.003,0.04] | 0.078 |
| Neuroticism | m6 | TAU | 397 | ACME | 0.004 [0.001,0.01] | 0.004 |
| | | | | ADE | 0.01 [-0.004,0.03] | 0.116 |
| | | | | Proportion | 0.24 [0.01,1.34] | 0.042 |
| | | | | Total | 0.02 [0.0006,0.04] | 0.038 |
| Neuroticism | m127 | TAU | 388 | ACME | 3.31E-3 [2.26E-4,7.22E-3] | 0.028 |
| | | | | ADE | 0.01 [-0.005,0.03] | 0.144 |
| | | | | Proportion | 0.01 [-0.005,0.03] | 0.144 |
| | | | | Total | 0.02 [-0.002,0.04] | 0.074 |
| Neuroticism | m131 | TAU | 388 | ACME | 0.002 [-5.88E-5,5.75E-3] | 0.06 |
| | | | | ADE | 0.01 [-0.004,0.04] | 0.144 |
| | | | | Proportion | 0.13 [-0.42,1.17] | 0.136 |
| | | | | Total | 0.02 [-0.002,0.04] | 0.08 |
| m7 | TAU | Cognitive decline | 388 | ACME | -0.41 [-0.63,-0.22] | <2.2E-16 |
| | | | | ADE | -0.51 [-0.9,-0.13] | 0.008 |
| | | | | Proportion | 0.45 [0.24,0.78] | <2.2E-16 |
| | | | | Total | -0.92 [-1.36,-0.51] | <2.2E-16 |
| m6 | TAU | Cognitive decline | 397 | ACME | -0.31 [-0.51,-0.13] | 0.002 |
| | | | | ADE | -0.41 [-0.74,-0.06] | 0.02 |
| | | | | Proportion | 0.43 [0.22,0.86] | 0.002 |
| | | | | Total | -0.72 [-1.1,-0.36] | <2.2E-16 |
| m127 | TAU | Cognitive decline | 388 | ACME | 0.22 [0.08,0.38] | 0.002 |
| | | | | ADE | 0.38 [0.08,0.71] | 0.016 |
| | | | | Proportion | 0.37 [0.14,0.74] | 0.002 |
| | | | | Total | 0.6 [0.28,0.97] | <2.2E-16 |
| m131 | TAU | Cognitive decline | 388 | ACME | 0.14 [0.03,0.28] | 0.01 |
| | | | | ADE | 0.3 [0.07,0.53] | 0.014 |
| | | | | Proportion | 0.32 [0.07,0.74] | 0.01 |
| | | | | Total | 0.44 [0.2,0.69] | <2.2E-16 |

*Effects are reported from causal mediation analyses using nonparametric bootstrapped confidence intervals (1000 iterations). The covariates adjusted in the model includes sex, age at death, PMI, RIN, experimental batch, study (ROS or MAP), education, and race.

Abbreviations: ACME, average causal mediation effect; ADE, average direct effect; TAU, abnormally phosphorylated Tau protein, AT8.

**Supplementary figure S1. The impact of other four personality traits on the overall cortical expressions of co-expressed genes**. The four personality traits are displayed in the order of **a** conscientiousness, **b** agreeableness, **c** openness and **d** extraversion. For each trait, the left panel shows the omnibus analysis and the right panel shows the comparisons of associations to cognitive decline. For the omnibus test on the left, each dot represents one module and its expected and observed associations with the personality traits (-log10 *P* values) are shown on X and Y axis, respectively. The expected *P* values assume a null distribution with no linear associations between the personality trait and expression level of co-expressed gene module after adjusting age at death, sex, education, race, postmortem interval, study (ROS or MAP), RNA-seq batch and RIN score. The grey and dark areas indicate the extreme ranges of the QQ plot as generated by chance at a threshold of *P*=0.05 and *P*=0.10, respectively. The 95% and 90% confidence intervals were empirically derived by randomly assigning participants with the score of the personality trait and repeating the analysis 1,000 times. Based on the distribution of the observed *P* values for all the 47 co-expressed gene modules, the overall association between the four personality traits and module expression was likely to have occurred by chance (*P*>0.05). For the right panel, each dot represents one module (the module number is listed next to each dot) and their previously reported *P* values for the cognitive decline (*Mostafavi 2018*) are shown on the Y axis and the *P* values (-log10 transformed) of the effect of neuroticism are shown on the X axis with the red dashed line for the threshold of *P*=0.05. The red dots are those that were previously reported to be associated with cognitive decline (*Mostafavi 2018*).

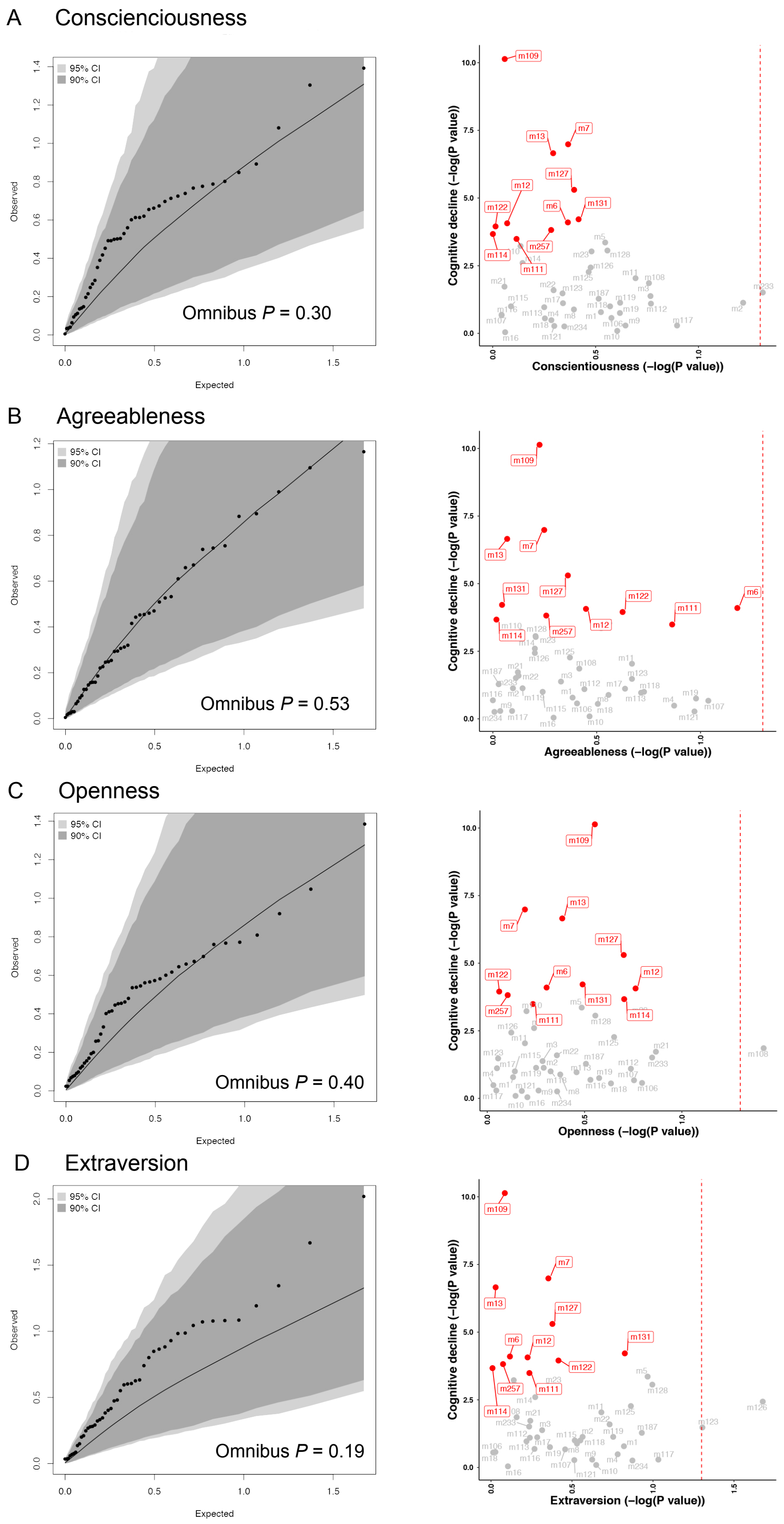

A Conscienciousness
95% CI
90% CI
Omnibus P = 0.30
Observed
Expected
Cognitive decline (−log(P value))
Conscientiousness (−log(P value))
B Agreeableness
95% CI
90% CI
Omnibus P = 0.53
Observed
Expected
Cognitive decline (−log(P value))
Agreeableness (−log(P value))
C Openness
95% CI
90% CI
Omnibus P = 0.40
Observed
Expected
Cognitive decline (−log(P value))
Openness (−log(P value))
D Extraversion
95% CI
90% CI
Omnibus P = 0.19
Observed
Expected
Cognitive decline (−log(P value))
Extraversion (−log(P value))

**Supplementary figure S2. Distribution of cortical module m6 and m131 expression in individuals with different levels of neuroticism**. Each dot represents one subject, and the subjects are distributed into 3 groups based on their neuroticism score: Group 1 with score from 0 to 12; Group 2 with score from 13 to 24; Group 3 with score from 25 to 36. On this scale, scores go from 0 (no neuroticism) to 36 (extensive neuroticism).

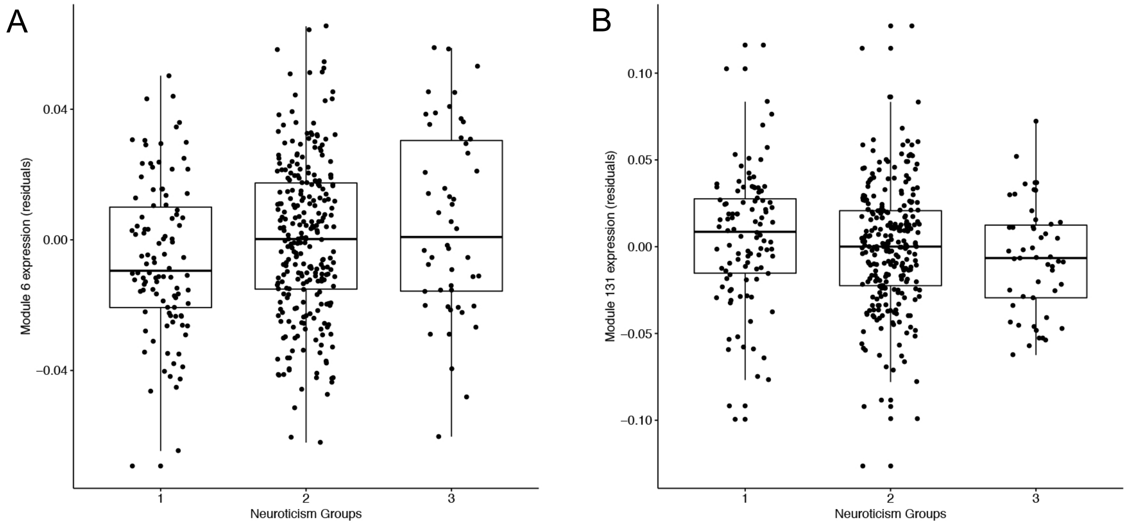
